# Detection of prions from spiked and free-ranging carnivore feces

**DOI:** 10.1101/2023.07.31.551307

**Authors:** H. N. Inzalaco, E. E. Brandell, S.P. Wilson, M. Hunsaker, D. R. Stahler, K. Woelfel, D. P. Walsh, T. Nordeen, D. J. Storm, S. S. Lichtenberg, W. C. Turner

## Abstract

Chronic wasting disease (CWD) is a highly contagious, fatal neurodegenerative disease caused by infectious prions (PrP^CWD^) affecting wild and captive cervids. Although experimental feeding studies have demonstrated prions in feces of crows (*Corvus brachyrhynchos*), coyotes (*Canis latrans*), and cougars (*Puma concolor*), the role of scavengers and predators in CWD epidemiology remains poorly understood. Here we applied the real-time quaking-induced conversion (RT-QuIC) assay to detect PrP^CWD^ in feces from cervid consumers, to advance surveillance approaches, which could be used to improve disease research and adaptive management of CWD. We assessed recovery and detection of PrP^CWD^ by experimental spiking of PrP^CWD^ into carnivore feces from 9 species sourced from CWD-free populations or captive facilities. We then applied this technique to detect PrP^CWD^ from feces of predators and scavengers in free-ranging populations. Our results demonstrate that spiked PrP^CWD^ is detectable from feces of free-ranging mammalian and avian carnivores using RT-QuIC. Results show that PrP^CWD^ acquired in natural settings is detectable in feces from free-ranging carnivores, and that PrP^CWD^ rates of detection in carnivore feces reflect relative prevalence estimates observed in the corresponding cervid populations. This study adapts an important diagnostic tool for CWD, allowing investigation of the epidemiology of CWD at the community-level.

## INTRODUCTION

The breakdown of detritus is important for regulating the movement of energy through food webs^1^, structuring ecological communities and ecosystem function^2^, and can also influence the dynamics of disease systems^3, 4^. Obligate and/or facultative scavenger species participate in scavenging, or the consumption of carrion, as a resource acquisition strategy. The latter are often also predators, consuming live prey. Scavenging and predation processes have important influences on disease epidemiology through removal of infectious individuals or materials that could transmit pathogens to new hosts^5, 6^. Simultaneously, scavengers and predators may also spread infectious diseases by transporting the pathogen to other areas (through consumption, subsequent fecal deposition, or translocation of contaminated material)^7^. Determining whether predators and scavengers reduce or enhance disease transmission risk in the host species they consume is important for understanding the ecological context of infectious diseases in wildlife. The role of these consumers in disease dynamics will be affected by factors such as carnivore diversity, gut capacity, digestive rates and fecal deposition patterns, pathogen environmental stability, host-pathogen encounters, and infectious dose.

Chronic wasting disease (CWD) is a highly contagious fatal neurological disease caused by a misfolded prion protein (PrP^CWD^) that affects several domestic and free-ranging cervid species. Currently, cervids are the only wildlife species monitored in routine CWD surveillance and the effects of other species and the environment on CWD dynamics are relatively unknown. For prion diseases, pathogen detection from abiotic and biotic environmental sources is technically challenging due to prions being devoid of genetic material. For years detection challenges limited the ability to address the sort of ecological questions readily possible for many other infectious agents that contain nucleic acids, such as many bacterial pathogens with complex disease ecology (e.g. *Bacillus anthracis*^8^, *Yersinia pestis*^9^, and *Francisella tularensis*^10^). Prion detection assays need to overcome the sensitivity limitations of existing techniques to be adaptable for detection of prions from a variety of ecological- and management-relevant environmental samples. As such, an assay with this capacity could be used not only to begin understanding how other species influence CWD processes but may also help to improve current surveillance efforts.

Although the first observations of CWD were made in captive mule deer (*Odocoileus hemionus*) and black-tailed deer (*Odocoileus hemionus columbianus*) in Colorado^11^, the disease has since spread broadly in cervid species across North America and Europe, including white-tailed deer (*Odocoileus virginianus*, hereafter referred to as WTD), elk (*Cervus canadensis*), reindeer (*Rangifer tarandus*), and moose (*Alces alces*)^12–15^. Through the disease course, PrP^CWD^ accumulates in body fluids and tissues^16, 17^ of presymptomatic and symptomatic CWD-infected individuals, is shed in excreta and secreta^18–21^, and ultimately results in neuroinvasion causing clinical disease and death^22, 23^. Transmission of CWD can occur either directly (i.e., contact between infectious and naive individuals) or indirectly through contact with contaminated environments^24^ Given that PrP^CWD^ can remain stable and infectious for years in the environment^25^, deposition and accumulation of PrP^CWD^ in the environment from infected carcasses or via shedding from infected individuals may lead to increased environmental disease exposure risk^26^. Consumption of CWD-infected hosts, through predation or scavenging suggests a role for carnivores in CWD epidemiology.

Coyotes (*Canis latrans*)^27^, American crows (*Corvus brachyrhynchos*)^28^, and cougars (*Puma concolor*)^29^ shed prions following ingestion of prion-infected material. Prion gut residency times in carnivores appear to be fairly short, lasting ≤ 3 days in coyotes and cougars, to just several hours in crows. Carnivores consuming contaminated cervid carrion may contribute to PrP^CWD^ translocation and contamination in the environment^27, 28^. However, infectious PrP^CWD^ shed in coyote feces was reduced relative to ingested material, implying feces-associated PrP^CWD^ deposits may contain less infectious material compared to unconsumed carrion^27^. Baune *et al.*^29^ suggest that most of the prions ingested by cougars are eliminated or sequestered, supporting the notion that predators may have a dilution effect on the CWD system. Further, a modeling study from Brandell *et al*.^5^ suggests that cougar and gray wolf (*Canis lupus*) predation pressure may independently decrease CWD outbreak size and delay prevalence increases of deer and elk, and this cleansing effect is amplified when predator selection for infected adults is greater than uninfected juveniles. Similarly, Fisher *et al*.^30^ recently reported predation by cougars may have slowed the increase in prevalence in an area of high CWD prevalence.

Previous studies investigating prions in scavengers relied on detection methods that range from mouse bioassays and antibody-based tools^28^, to an ultra-sensitive *in vitro* protein amplification assay, protein misfolding cyclic amplification (PMCA)^27^. Although mouse bioassay remains the gold standard for definitively assessing the presence and titer of PrP^CWD^, these assays can take months to years to complete. Bioassays are also often used in conjunction with PMCA, which offers the same level of sensitivity as mouse bioassay and is far more sensitive than antibody-based detection tools^31, 32^. For example, in the study by Nichols *et al*.^27^, PMCA was used to characterize the fecal samples prior to use as inoculum in the mouse bioassay. However, PMCA is costly and not adaptable for high-throughput diagnostics as it still requires the need for prion disease susceptible brain tissue as a conversion substrate and can take up to two weeks to produce a result. The development and application of an alternative ultrasensitive *in vitro* protein amplification assay, the real-time quaking-induced conversion (RT-QuIC), has advanced high throughput detection of PrP^CWD^ from host tissues^33, 34^ and various host secreta and excreta^19, 34–38^ without the use of animal tissue substrates. Currently, there has been only one report of the application of RT-QuIC for PrP^CWD^ detection from feces of cervid consumers (cougar), based on captive animals in a controlled laboratory environment^29^.

Here we evaluated the utility of RT-QuIC for detection of PrP^CWD^ using fecal-spiking assays from a suite of relevant predator and scavenger species. We assessed limits of PrP^CWD^ recovery and detection sensitivity from nine species relative to RT-QuIC characterized CWD-positive brain tissue. We then used this assay to detect PrP^CWD^ in feces samples from two free-ranging carnivore species – coyotes and cougars. Our findings from field collected feces suggests that this approach could be used to institute early surveillance of CWD, especially in locations or jurisdictions neighboring CWD endemic zones that are considered areas of concern for geographic spread or areas where hunter-harvest rates are low, and also to begin to unravel how other species influence CWD dynamics and geographic spread.

## MATERIALS AND METHODS

### Fecal sample spiking assays

Individual fecal samples were handled separately, using fresh disposable, single use nitrile gloves, and disposable weigh boats to prevent cross-contamination of samples. Using CWD-negative fecal samples as determined by RT-QuIC described below data analysis, spiking experiments were used to demonstrate recovery of PrP^CWD^ from feces of 9 different scavenger and predator species: gray wolf (hereafter wolf), cougar, coyote, American crow (hereafter crow), American black bear (*Ursus americanus*, hereafter bear), raccoon (*Procyon lotor*), common raven (*Corvus corax*, hereafter raven), red fox (*Vulpes vulpes*, hereafter fox), and bald eagle (*Haliaeetus leucocephalus*, hereafter eagle; sources listed above and in Table 1). Spiking assays were carried out by using 50 mg of each fecal sample combined with 50 μL of a 10-fold dilution of a 10% (w/v) CWD-positive WTD brain homogenate (BH) from the obex region of tagged WTD #5219 in the late stages of CWD (hereafter referred to as the reference sample), provided by the Wisconsin Department of Natural Resources (WDNR). The reference sample was prepared in 1X phosphate-buffered saline (PBS) to achieve a range of final concentrations of 10^-3^ to 10^-8^ mg/mL of CWD-positive brain. CWD-positive and negative BH were determined by RT-QuIC. For the negative control spike, 50 μL of a 10^-3^ mg/mL dilution of a 10% (w/v) CWD-negative WTD BH was used. Each spike dilution was added to feces and allowed to soak in for 24 hours, then placed in a 50 °C incubator without agitation for 16 h, to keep assay temperature conditions consistent with those used for RT-QuIC. Spiked fecal samples were then prepared as described below.

**Table 1.** Species fecal sample collection location information, use, and storage conditions for.

| Species | Number of<br>samples tested in<br>triplicate | Year collected | Location of collection |
| --- | --- | --- | --- |
| **Fecal samples used for RT-QulC spiking assays** |  |  |  |
| Wolf | 1 | 2020 | Northern Yellowstone Ecosystem |
| Coyote | 1 | 2022 | Northern Yellowstone Ecosystem |
| Fox | 1 | 2022 | Northern Yellowstone Ecosystem |
| Raven | 1 | 2022 | Northern Yellowstone Ecosystem |
| Cougar | 1 | 2022 | Northern Yellowstone Ecosystem |
| Bear | 1 | 2022 | Minnesota Wild and Free Wildlife<br>Rehabilitation Center (MWF) in Garrison,<br>Minnesota. |
| Raccoon | 1 | 2022 | MWF |
| Eagle | 1 | 2022 | MWF |
| Crow | 1 | 2022 | Dane County Humane Society Wildlife<br>Center, Madison, Wisconsin, U.S.A. |
| **Fecal samples used for RT-QulC diagnostics assays** |  |  |  |
| Coyote | 6 | 2021-2022 | Iowa County, Wisconsin, U.S.A. |
| Cougar | 5 | 2012 & 2014 | Niobrara Valley, Cherry County, Nebraska,<br>U.S.A. |
| Cougar | 10 | 2010 & 2021 | Pine Ridge region, Dawes and Sheridan<br>Counties, Nebraska, U.S.A. |

samples included in this study. All samples were stored at $-80^{\circ}\text{C}$ .

### Fecal sample extract preparation for RT-QuIC assay

Fecal samples (Table 1) were stored at −80°C prior to preparation for use in the RT-QuIC assay. Individual fecal samples and subsamples were handled separately, using fresh disposable/single use nitrile gloves, and disposable weigh boats to prevent cross-contamination of samples. To prepare field collected fecal samples for detection of PrP^CWD^, each individual sample had one subsample collected from three different areas/regions consisting of a slice from the middle and both ends of the sample, totaling three biological replicates (50 mg each). A separate scalpel blade was used to cut each section, that included surface material as well as inner material up to a depth of 1 cm to ensure sufficient sampling of surface and inner portions. These 3 subsamples were then extracted individually, for a total of 3 separate extractions, using 1 mL of sterile 1X PBS, then processed in a thermomixer (1,400 rpm, 30 min, 25°C; Eppendorf ThermoMixer F1.5) followed by centrifugation at 16,000 x *g* for 15 min. Supernatants (∼750 μL) were collected, followed by addition of 750 μL 1X PBS to each feces pellet and remaining buffer. Samples were vortexed, thermomixed (1,400 rpm, 30 min, 25°C), and centrifuged again at 16,000 x *g* for 15 min, after which an additional ∼750 μL supernatant was removed and added to the first supernatant. Resultant supernatants were then centrifuged again for further clarification at 16,000 x *g* for 15 min, followed by collection of 1 mL of clarified supernatant into a new sterile tube and 500 μL of 23.1 mM sodium phosphotungstate hydrate (Na-PTA; Sigma-Aldrich, Cat. # 496626) was added. Samples were incubated without agitation for 16 hours at 4°C, then centrifuged (4 °C, 30 min, 5000 x *g*). Sample supernatants were carefully removed from each tube and discarded. Pellets were retained and washed with a 1:1 solution of 18 MΩ H_2_O and 23.1 mM Na-PTA followed by centrifugation (4 °C, 30 min, 5000 x *g*) and aspiration of the wash solution. Pellets were resuspended in 30 µL of RT-QuIC sample buffer (0.1 g/mL sodium dodecyl sulfate in phosphate-buffered saline with N-2 cell culture supplement; ThermoFisher, Waltham, MA) and reconstituted using a Qsonica Q700 cup horn ultrasonicator (Amplitude 36 for 1 min.). A volume of 2 µL of each reconstituted sample was used to seed each reaction well for 8 technical replicates.

### RT-QuIC Assay

The RT-QuIC *in vitro* prion amplification assay was performed as described by Orru *et al*.^39^, with sodium iodide as described by Metrick *et al*.^40^ with minor modifications. Briefly, 2 µL of sample extracts were added to a given well of a 96-well format optical-bottom black microplate (Thermo Scientific, Fair Lawn, NJ, USA), each already containing 98 µL of RT-QuIC reaction mixture (0.1 mg·mL^-1^ 90-231 recombinant hamster prion protein, produced as previously described:^39^, 300 mM sodium iodide, 20 mM sodium phosphate, 1.0 mM ethylenediaminetetraacetic acid, and 10 µM thioflavin T). Microplate-compatible spectrophotometers capable of heating, shaking, and fluorescence monitoring (BMG FLUOstar, Cary, North Carolina) were used with the following instrument settings: 50 °C for spiked samples double orbital pattern shaking at 700 rpm with 60-s shake / 60-s rest cycles, fluorescent scans (λ_excitation_ = 448 nm, λ_emission_ = 482 nm) every 15 minutes, at a gain of 1,600, and a total run time of 48 h.

### Collection of fecal samples

Fecal samples included in this study were obtained from either the field or from wildlife rehabilitation centers (Table 1). Individual fecal samples were collected using sterile Whirl-Pak bags and sterile, single-use nitrile gloves (changed between sample collects), then stored at −80 °C prior to analysis. Below we describe location information and methods used to locate and collect fecal samples.

#### Northern Yellowstone ecosystem

Wolf, cougar, fox, coyote, and raven fecal samples included in this study from the Northern Yellowstone ecosystem (NYE) were provided by the Yellowstone Center for Resources, Yellowstone National Park, Wyoming, U.S.A. The NYE is a mountainous region (ranging approximately 1500–2400 m) located in northwestern Wyoming, USA, and south-central Montana, USA and is characterized primarily by lower elevation steppe and grassland, and higher elevation coniferous forests, with relatively few wetland areas^41^. The NYE is inhabited by several cervid prey species on which wolves and other predators prey^42^. CWD had not been detected in the study area during sampling periods, nor had CWD been detected in cervids within 30 kilometers of the study area. Fecal collection methods for wolf^43^, followed by fox, coyote, cougar, and raven from the NYE are described briefly. Feces from GPS radio collared wolves (IACUC IMR YELL Smith wolves 2012) were collected in 2020 during early winter (30 days: November 15-December 15) approximately 17 days following deposition. Cougar, coyote, and fox fecal samples were located and collected from cougar GPS clusters near cougar kill sites or by tracking fox and coyotes opportunistically during cougar cluster investigations. Raven fecal samples were collected during capture, tagging, and handling of ravens for a monitoring study following permits and animal handling protocols permitted by state (Montana Scientific Collector’s Permit 2022-020-W), federal (Master Banding Permit 22489), NPS (Yell-2022-SCI-8072), and University of Washington animal care and use committee (Protocol 3077-01). All methods for feces collection were performed in accordance with the relevant guidelines and regulations.

#### Southwest Wisconsin

As part of a multiyear CWD study centered in southwest Wisconsin, USA the Wisconsin Department of Natural Resources (WDNR) captured and fit 763 deer with GPS collars from 2017 to 2020 in Iowa, Grant, and Dane counties in southwestern Wisconsin, following protocols approved by WDNR’s Animal Care and Use Committee (Protocol: 16-Storm-01). This region is characterized by high CWD prevalence^44^— an area where CWD was first established in WI, has been estimated to have been present in the environment for over 20 years^45^, and studied extensively in prior research^46–48^. Habitat in this area is characterized by steep hills, forested ridges, deep river valleys, karst geology, and cold-water trout streams. Elevations range from 184 to 524 m. Coyote are reported to leave numerous fecal deposits within 80 meters of carrion they are consuming^49^, therefore fecal samples were collected by the field crew opportunistically within 0-80 meters of collared WTD mortality sites from 2021-2022. Deer GPS collars notified the field crew when the collar had been motionless for 4 or more hours. Deer mortality sites were then identified by GPS points and/or VHF telemetry. Morality sites were investigated to determine cause of death and those that had signs of predator activity were searched for feces.

#### Northern Nebraska

The cougar fecal samples from the Pine Ridge and Niobrara Valley regions in northern Nebraska, U.S.A. used in this study were provided by Nebraska Game and Parks Commission (NGPC). Habitat in the Rine Ridge area is characterized by meadows, pine and deciduous forests, steep buttes, small streams, and minor peaks (ranging approximately 900-1600 m). Habitat in the Niobrara Valley is characterized by steep hills, bluffs, pine forests and canyons, boreal forests, grasslands, and the Niobrara River. As part of ongoing carnivore studies in Cherry, Dawes, and Sheridan counties in Nebraska, U.S.A., cougar fecal samples were collected by NGPC wildlife staff with the help of a trained detection dog and handler. While not part of ongoing CWD surveillance efforts in Northwest Nebraska, cougar fecal samples included in this study were collected within areas that are also designated big game research deer management units (DMUs) where CWD has been detected.

#### Minnesota Wildlife Rehabilitation Center

Feces from adult bear, raccoon, and eagle were collected during the month of September 2022 at the Minnesota Wild and Free Wildlife Rehabilitation Center (MWF) in Garrison, Minnesota, USA. Animals that fecal samples were collected from were originally found at different geographic locations prior to being brought to WMF, varied in time post admittance to MWF, and had differing diets as appropriate for each species. An admitted bear was found in Polk County, Minnesota, USA and the admitted raccoon and eagle were found in Cass County, Minnesota, USA. Individuals whose feces were included in this study were admitted between the months of May-July and were transported to MWF by The Minnesota Department of Natural Resources (MNDNR). The bear diet consisted of dry dog food supplemented with produce (i.e., apples, melon, corn, and assorted berries). The eagle diet typically included fish and rodents (i.e., chipmunks, mice, or gophers). The raccoon diet consisted of dry and/or wet dog food mixed with assorted produce (i.e., apples, melon, corn, and assorted berries).

#### Dane County Humane Society Wildlife Center

Feces from a juvenile crow was collected during the month of May 2022 at the Dane County Humane Society Wildlife Center in Madison, Wisconsin, USA. The crow was admitted in early May and was fed a diet that consisted of eggs, chicken, meal worms, seeds, nuts, and produce (i.e., apples, melon, corn, and assorted berries).

### Data Analysis

Amyloid formation rate (AFR) data generated from the RT-QuIC assays were analyzed and visualized using Jmp Pro 15 (SAS Institute, Cary, NC) and Prism 8 (GraphPad, San Diego, CA). To apply a rigorous standard for distinguishing true positive samples from true negatives, the AFR threshold times (i.e., the time at which amplification is determined to have occurred in the RT-QuIC assay) were calculated by adding twenty times the standard deviation of the relative fluorescence unit (RFU) values from cycles 3-14 to the mean of RFU values from cycles 3-14. We previously applied this method to account for baseline fluorescence variation amongst samples in determining if the sample was PrP^CWD^ positive^50^. Due to false seeding observed in crow, bear, eagle, and raccoon negative control fecal samples, additional analysis was required to distinguish true seeding from false seeding events for these species. To accomplish this, empirical distributions of threshold times in hours of 24 replicates of unspiked fecal samples were used to determine a threshold time that would yield a specificity ≥ 95% for fecal samples from each species. These data were then used to determine a cycle end-time that excluded ≥95% of the false seeding that occurred for each species. Cycle end-times for these four feces were determined to be 20 h for crow (specificity of 95%), 24 h for bear (specificity 100%), 25 h for eagle (specificity 96%), and 17 h for raccoon (specificity 100%) (Supplementary Fig. S1). All fecal samples analyzed by RT-QuIC were considered positive if a sample had at least 3 out of 8 technical replicates (or half or more of 24 replicates) with seeding activity and had AFR values that were significantly different from control AFR values based on statistical analysis. Analyses for spiked samples were assessed with Dunnett’s multiple comparison tests, while for the surveillance samples, Kruskal-Wallis tests was used to distinguish which samples had AFRs significantly different from the northern Yellowstone ecosystem (NYE) negative control fecal sample.

Using AFR as a relative measure for prion concentration in a sample, we first evaluated our ability to detect PrP^CWD^ from spiked fecal samples compared to the reference sample, and if the AFR differed among carnivore species. We used a two-way (factorial) ANOVA to compare AFR values between sample types (CWD-positive brain or spiked fecal samples from 9 different species). We included an interaction between species and spike dilution; to assess if detection/recovery in the different feces was sensitive to the concentration of the spike across the 10-fold dilution series. Significant interactions were determined using the Tukey HSD for multiple comparisons.

Amyloid formation rates for field collected coyote and cougar fecal samples were found to be non-normally distributed (Supplementary Figure S2). As such, the nonparametric Kruskal-Wallis test was applied to compare individual fecal samples. If statistical differences were observed across each individual fecal sample, then the non-parametric Steel-dewass post hoc test was used to determine which individual samples differed from the negative control fecal sample.

## RESULTS

Using RT-QuIC, our experimental spiking studies aimed to test extraction efficiency and evaluate the presence of assay inhibitors through detection sensitivity of PrP^CWD^ in feces from different scavenger and predator species using the reference sample as the source material for spiking in all experiments (Fig. 1, see Table 1 for feces source information. Fecal samples for each species were spiked with 10-fold dilutions of brain-derived PrP^CWD^ (Fig. 1). Variation in assay sensitivity for the different dilutions of BH spike was observed across species. The limit of relative PrP^CWD^ detection was significantly different from the negative controls in a species-specific fashion. The most sensitive detection was from wolf feces, with 8/8 technical replicates showing seeding activity (defined as having relative fluorescent units above the RFU threshold) in feces spiked with only 10 ng of CWD-positive WTD BH. This was followed by eagle (6/8 seeding), which had a detection limit at 100 ng of brain-derived PrP^CWD^ (Fig. 1). Detection limits were higher for other species, at a 10^3^ ng spike for crow (6/8), bear (3/8), cougar (4/8), and raven (8/8), a 10^4^ ng spike for coyote (8/8) and raccoon (4/8) (Fig. 1), and a 10^5^ ng spike for fox (8/8) (Fig. 1).

**Figure 1.**
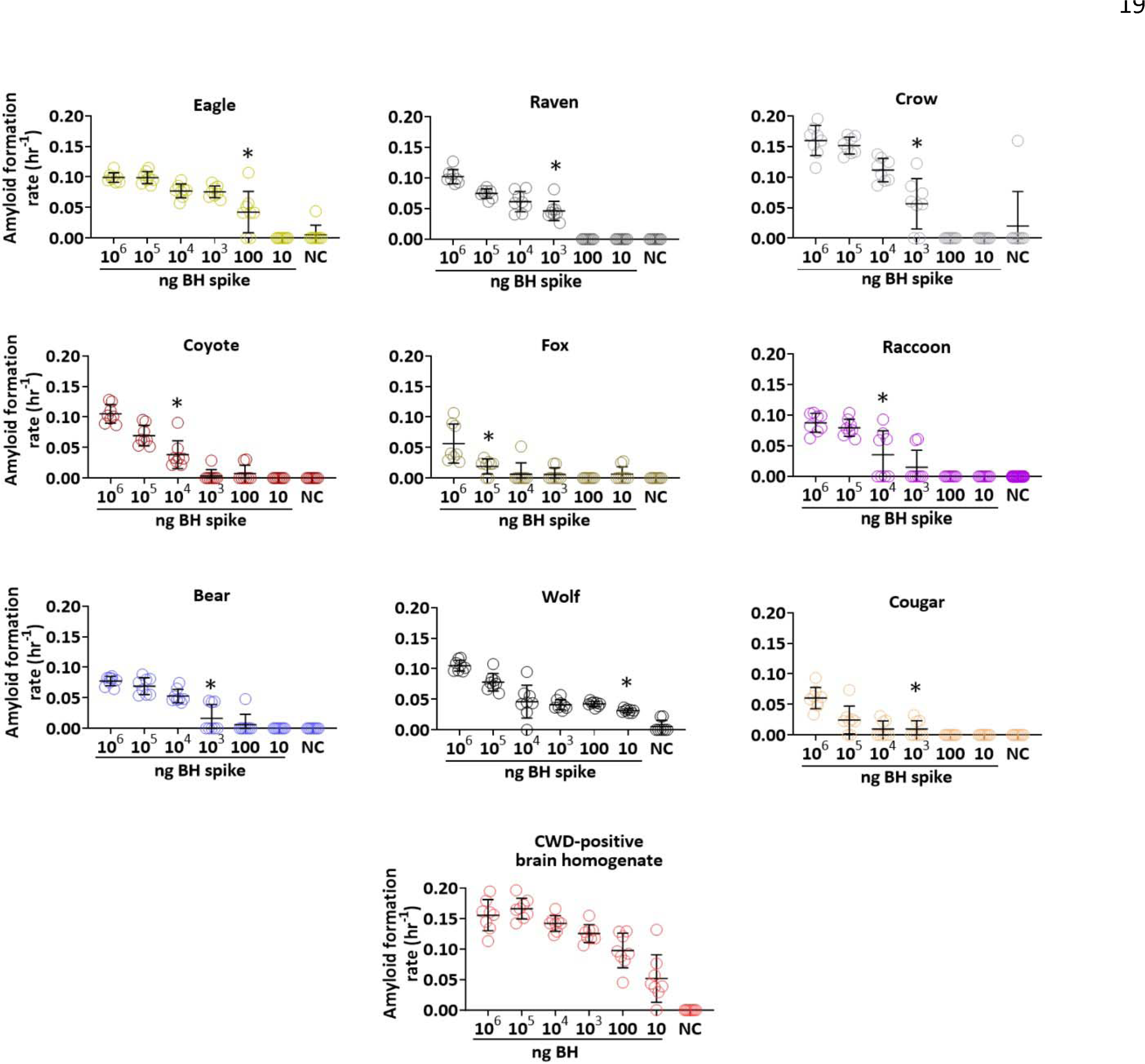
Recovery and detection sensitivity of chronic wasting disease (CWD) prions (PrP^CWD^) from spiking experiments of predator and scavenger feces by the real-time quaking-induced conversion (RT-QuIC) assay. Amyloid formation rates (AFRs) by RT-QuIC for predator and scavenger fecal samples spiked with 10-fold dilutions of 10% brain homogenate (from the obex region; BH) shown in nanograms (ng) from a CWD-positive white-tailed deer. Data points of AFRs ± standard deviation of eight technical replicates across each spiking dilution series in feces samples from crow, raven, eagle, raccoon, grey wolf, coyote, fox, cougar, and bear are shown. The dilution curve for the BH used as spiking material is shown for comparison to PrP^CWD^ recovery and detection sensitivity in feces. * = limit of detection sensitivity of spiked material in each species. NC = negative control (feces without a BH spike).

The variable detection limits observed among species may be due to differences in inhibition of the assay by the fecal material, the extraction process, or a combination of both. Overall, we found differences across species do account for some of the variation seen in the magnitude of AFRs for spiked feces when compared to dilutions of the reference sample on its own (R^2^ = 0.90, *F*_9, 420_ = 155.46, *p* < 0.0001). Mean AFRs for all species were significantly lower compared to pure CWD-positive WTD BH and there were species differences in PrP^CWD^ recovery (Tukey HSD post-hoc test for multiple comparisons, Supplementary Table S1, Fig. 2a). Spiked crow, eagle, and wolf feces had significantly higher AFRs than the other species (Supplementary Table S1, Fig. 2a). Cougar and fox feces had the lowest mean AFRs (Supplementary Table S1, Fig. 2a). AFR values from spiked raven did not differ significantly from wolf, bear, coyote, or raccoon feces, however AFR values of bear, coyote, and raccoon fecal samples did differ from the other species (Supplementary Table S1, Fig. 2a). AFRs for spiked fecal samples from all species decreased across the dilution series (*F*_5,420_ = 355.69, *p* < 0.0001), with a significant interaction between feces and spike dilution (*F*_45, 420_) = 9.36, *p* < 0.0001). The magnitude of differences in observed AFRs varied among species and by spike dilution, however there was a general humped-shaped pattern observed for differences in recovery of PrP^CWD^ across many of the species (Fig. 2b). The largest differences in recovery were observed at mid-level dilutions rather than for the lowest and highest dilutions for all spiked feces samples, with an exception for eagle and crow (Tukey HSD post-hoc tests, Fig. 2b, Supplementary Table S2).

**Figure 2.**
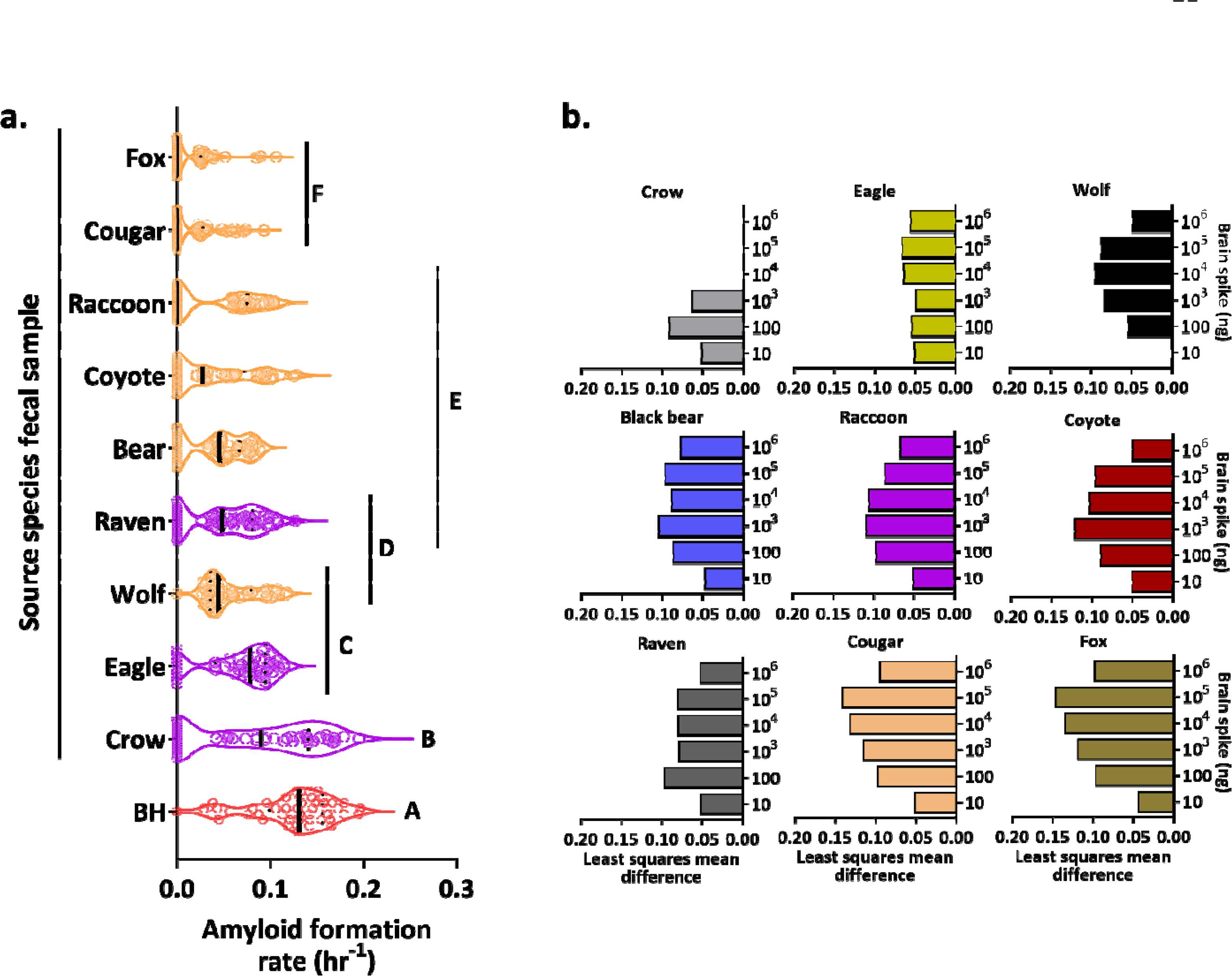
Differences in recovery and detection sensitivity of chronic wasting disease (CWD) prions (PrP^CWD^) from spiked carnivore feces. (a) Distribution of amyloid formations rates (AFRs) across spiked fecal samples showing the median (solid black line) and first and third quartiles (dotted black lines). CWD-positive brain homogenate (BH, the reference sample) is shown in red, mammalian species in orange and avian species in purple. Letters show which species are statistically different by Tukey HSD post-hoc test. **(b)** Illustration of the significant least squares mean AFR value differences between 10-fold dilutions of CWD-positive brain homogenate and recovered brain-derived PrP^CWD^ from spiked fecal samples by species and brain spike dilutions using the Tukey HSD post-hoc test. For dilutions lacking a bar in the plots, there was no significant difference in AFR between fecal samples and the reference sample.

Following these proof-of-concept spiking experiments, we then determined if we could detect PrP^CWD^ in feces from free-ranging predators/scavengers from areas with CWD. We examined coyote and cougar fecal samples collected from three distinct geographic regions where CWD has been detected. In Iowa County, Wisconsin, six different coyote fecal samples were collected within 80 meters of mortality sites for six different collared WTD in an area where CWD prevalence is ∼40% and 30% in adult males and females, respectively^51^. Of these WTD carcasses, 5/6 were confirmed CWD-positive by RT-QuIC on tissues (i.e., brain or skin from ear or belly; Fig. 3).

**Figure 3.**
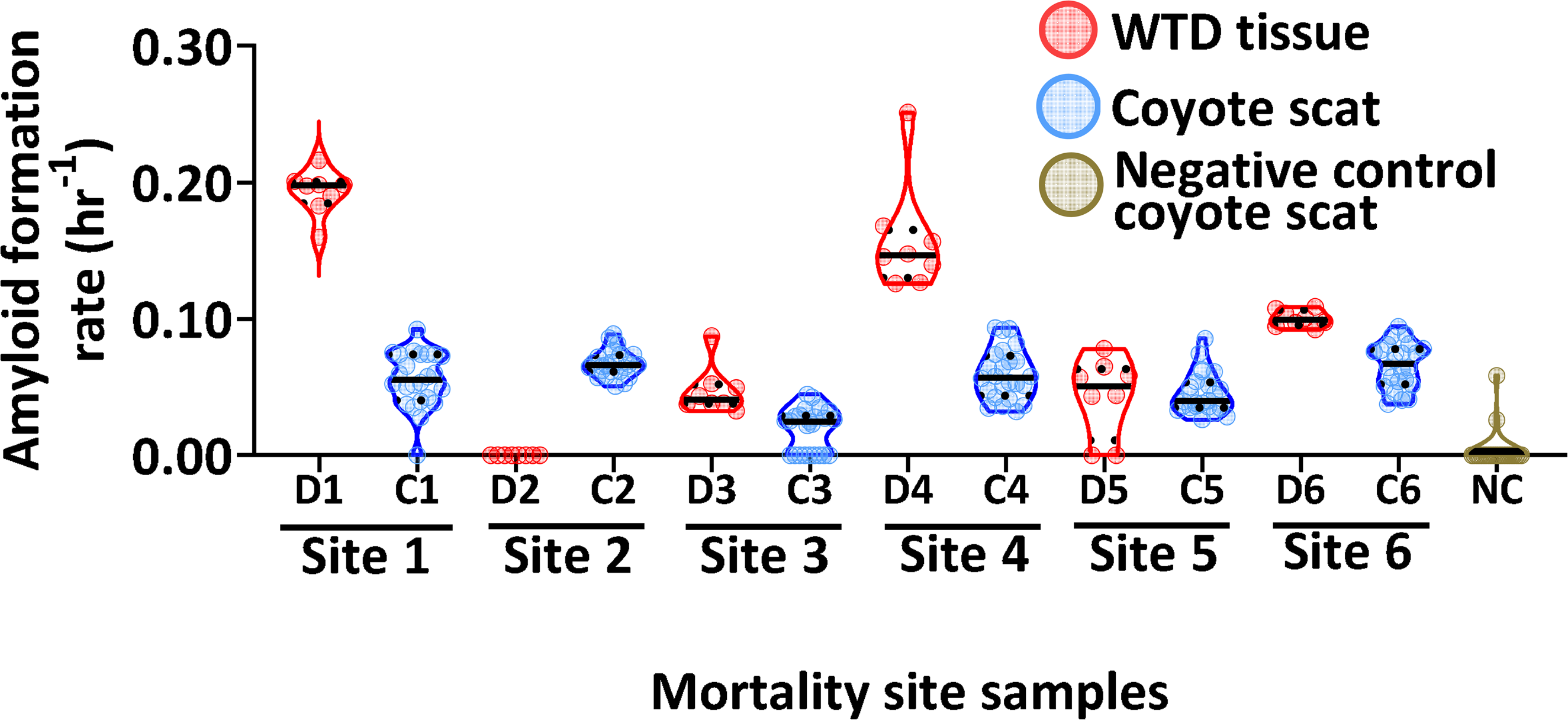
The presence of chronic wasting disease (CWD) prions (PrP^CWD^) in coyote fecal samples collected near collared white-tailed deer (WTD) mortality sites within Iowa County, Wisconsin, USA. Comparisons of real-time quaking-induced conversation (RT-QuIC) amyloid formation rates (AFR) of tissues (D1, obex; D2, obex; D3, belly skin; D4, ear pinna; D5, obex; D6, skin) from six different WTD carcasses (red) with six different coyote fecal samples found near each respective mortality site (blue). Samples are grouped by WTD mortality site ID (by number; D=deer and C=coyote from each site) on the x-axis. Data points show 8 and 24 replicates (3 separate extractions, 8 technical replicates each) of WTD tissue or coyote feces, respectively, with the median (solid black line) and first and third quartiles (dotted black lines). Negative control (NC; tan) represents 3 separate extraction results of 8 technical replicates each for coyote feces from 1 individual collected from the Northern Yellowstone Ecosystem.

We also evaluated how variable PrP^CWD^ presence is across a single sample and how repeatable our results for RT-QuIC were across plates by sampling and separately extracting and testing three different regions of a given sample. RT-QuIC assays of three separate subsamples from each coyote fecal sample (labeled as C1-C6) showed seeding activity in 8/8 of subsamples, with exception of sample C3, where all eight technical replicates for the first and third extractions showed seeding activity but had no seeding activity for the second extraction (Table 2). Coyote feces AFRs were significantly greater than the controls (Kruskal-Wallis test: Χ^2^ = 106.29, p<0.0001, df = 6; Steel-dwass post hoc test: *p*≤ 0.00016; Fig. 3; Table 2). Amyloid formation rates of coyote feces were lower than those of WTD tissue from each mortality site but had similar value ranges across each individual fecal sample and very little variation among technical replicates of each scat subsample (Fig. 3).

**Table 2.** Presence or absence of chronic wasting disease (CWD) prions (PrP^CWD^) from subsamples of coyote (*Canis latrans*) fecal samples collected from within Iowa County, Wisconsin, USA and one negative control (NC) coyote fecal sample collected from the Northern Yellowstone Ecosystem (this separates the biological replicates shown together in Figure 3, to note repeatability across subsamples) assessed by the real-time quaking-induced conversion (RT-QuIC) assay. Values indicate the proportion of seeding activity for three separate extractions, 8 technical replicates each (24 total replicates) for each fecal sample.

| Sample ID | Extraction 1 | Extraction 2 | Extraction 3 | Total seeded reactions | PrP<sup>CWD</sup><br>+/- for each<br>extraction |
| --- | --- | --- | --- | --- | --- |
| C1 | 8/8 | 7/8 | 8/8 | 23/24 | +/+/+ |
| C2 | 8/8 | 8/8 | 8/8 | 24/24 | +/+/+ |
| C3 | 7/8 | 0/8 | 8/8 | 15/24 | +/-/+ |
| C4 | 8/8 | 8/8 | 8/8 | 24/24 | +/+/+ |
| C5 | 8/8 | 8/8 | 8/8 | 24/24 | +/+/+ |
| C6 | 8/8 | 8/8 | 8/8 | 24/24 | +/+/+ |
| NC | 0/8 | 1/8 | 1/8 | 2/24 | -/-/- |

Cougar fecal samples were collected from DMUs in the Niobrara Valley (2012: n=2; 2014: n=3) and Pine Ridge area (2010: n=5; 2021: n=5) in Cherry, Dawes, and Sheridan Counties, Nebraska, USA. AFRs of three Niobrara (one from 2012, and two from 2014) and three Pine Ridge 2021 samples were significantly greater than the controls (Kruskal-Wallis test: Χ^2^ = 255.25, p<0.0001, df = 15; Steel-dwass post hoc test: *p*≤ 0.0226; Fig. 4). RT-QuIC assays of three separate subsamples taken from each fecal sample showed variable seeding activity within the 8 technical replicates, ranging from 2/8 to 8/8 (Table 3). Out of 24 replicates, at least half had seeding activity for each of these six fecal samples (Table 3). The range of AFRs among these fecal samples were variable, where values from the Pine Ridge 2021 samples were most elevated compared to the three samples from Niobrara (Fig. 4). AFRs for the remaining samples were not significantly greater than the controls (Steel-dwass post hoc test: *p*≥ 0.05; Fig. 4). RT-QuIC assays of three separate subsamples taken from each of these individual fecal samples also showed variable seeding that ranged from 0/8 to 6/8 (Table 2). The total number of replicates (out of 24) with seeding for each of these fecal samples was less than half (Table 3). Overall, subsample replicate AFRs from cougar fecal samples (Fig. 4, Table 3) were more variable than those seen from coyote feces (Fig. 3, Table 2).

**Figure 4.**
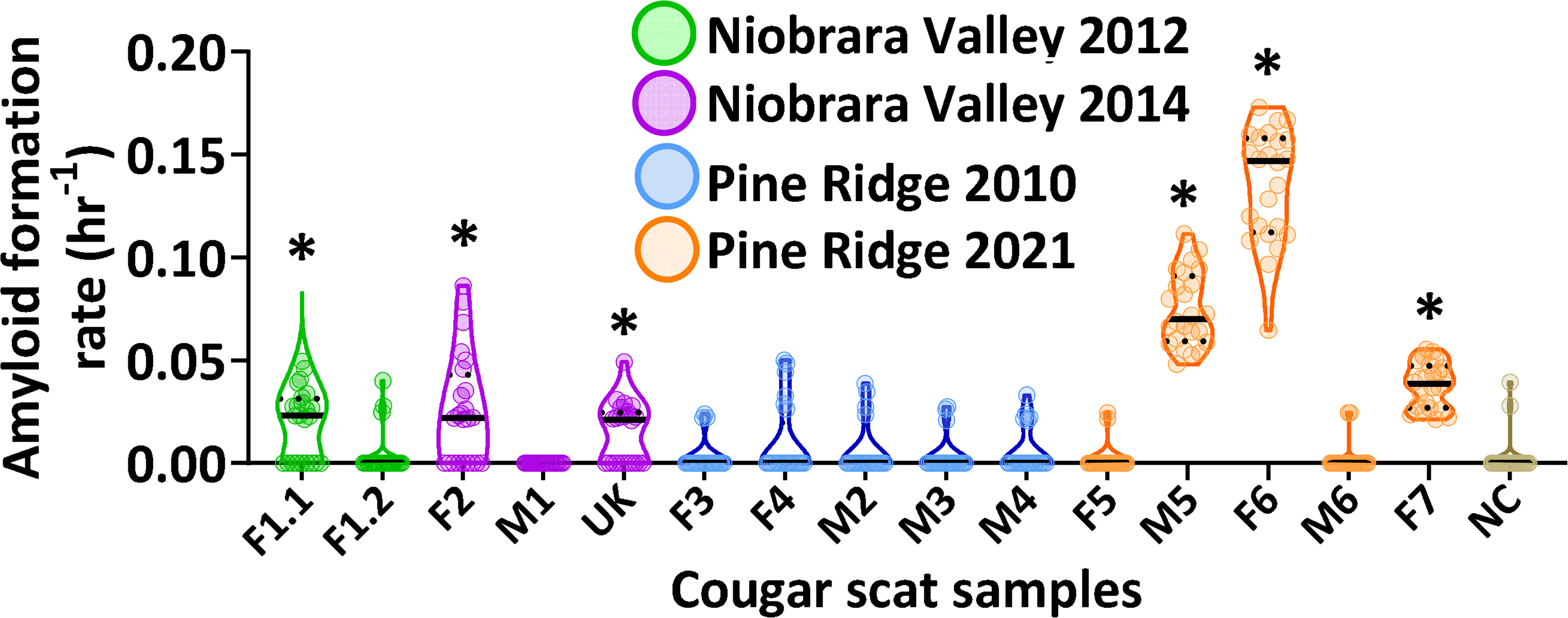
Presence of chronic wasting disease (CWD) prions (PrP^CWD^) in cougar fecal samples collected from two locations within Cherry, Dawes, and Sheridan Counties, Nebraska, USA. Real-time quaking-induced conversion (RT-QuIC) amyloid formation rates (AFR) from cougar feces collected from different individuals from the Niobrara Valley in 2012 (green) and 2014 (purple) and in 2010 (blue) and 2021 (orange) from the Pine Ridge region. Samples are grouped by individual cougar ID on the x-axis (F1.1 and F1.2 are two separate samples from the same individual). Data points shown are for 24 technical replicates (3 separate extractions of 8 technical replicates each), with the median (solid black line) and first and third quartiles (dotted black lines). Negative control (NC; tan) represents triplicate assay results for CWD-negative cougar feces from 1 individual collected from the Northern Yellowstone Ecosystem. * = positive for CWD based on Steel-dwass post-hoc test (*p* ≤ 0.05).

**Table 3.**
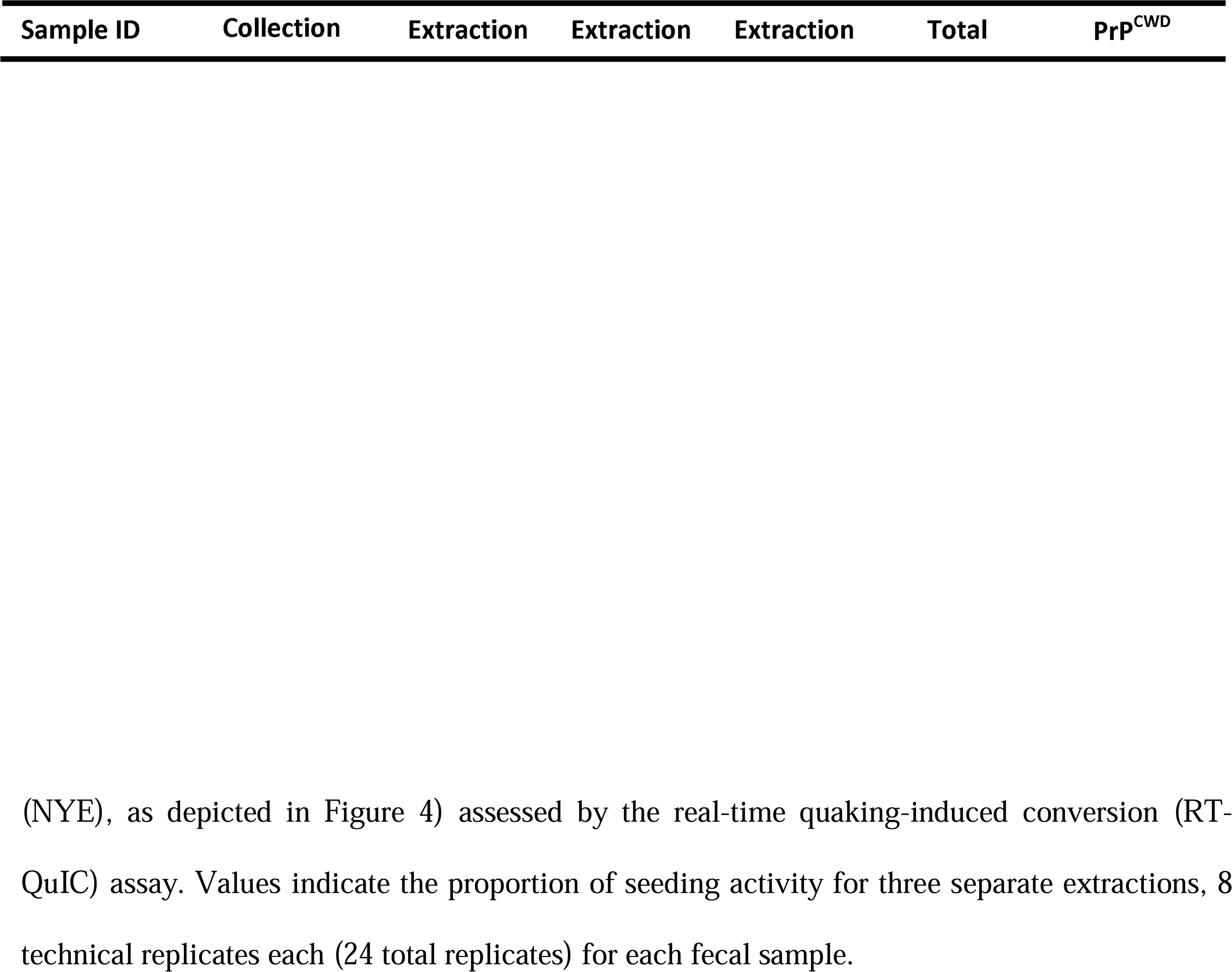
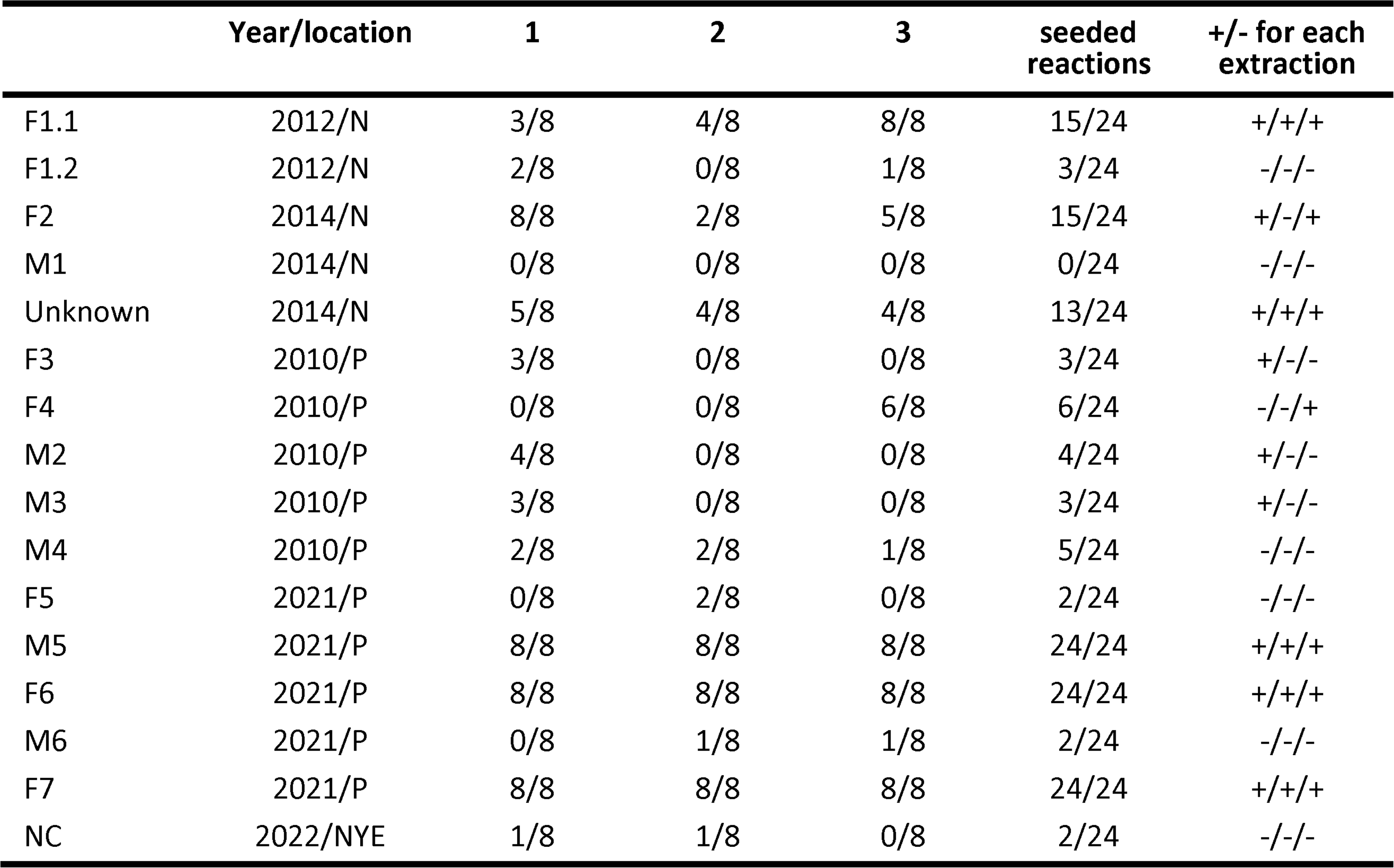
Presence or absence of chronic wasting disease (CWD) prions (PrP^CWD^) for fecal samples from 15 different cougars (*Puma concolor*) from the Niobrara (N) of Pine Ridge (P) deer management units within Cherry, Dawes, and Sheridan Counties, Nebraska, USA and one negative control (NC) cougar fecal sample (collected from the Northern Yellowstone Ecosystem.

Results of PrP^CWD^ occurrence in cougar feces from the 2010 (0/5) and 2021 (3/5) Pine Ridge samples increase from 2010 to 2021 within this DMU^52^ (Fig. 4, Table 3). Prevalence of CWD in WTD for the Pine Ridge DMU was 7% (21/298; SE= 0.015) in 2010. Although host CWD surveillance was not conducted in 2021 for the Pine Ridge DMU, surveillance in 2019 and 2022 estimated WTD CWD prevalence of 26% (69/264; SE= 0.027) and 27% (62/244; SE= 0.028), respectively. The DMU where the Niobrara Valley cougar feces were collected has had host CWD prevalence estimates of under 1% for the past 18 years. Estimated prevalence in 2012 was 0% (0/512), and although no surveillance was conducted from 2013-2016 for this DMU, 2018 prevalence was reported to be, 0.41% (1/243; SE= 0.004)^52^. Although a direct comparison of PrP^CWD^ detection between cougar scat and WTD is difficult to make due to prevalence estimates being very sensitive to small sample sizes^53^, it appears that PrP^CWD^ can be detected in carnivore scat even when CWD prevalence in WTD is very low.

## DISCUSSION

The role of sympatric wildlife in CWD epidemiology remains incompletely understood, but cervid-consuming predators and scavengers may alter rates of disease spread through removal or dispersal of PrP^CWD^ on the landscape. Determining the effects of cervid consumers on CWD ecology and epidemiology could be facilitated with development of non-invasive and sensitive methods with high throughput capacity for PrP^CWD^ detection. We presented results from laboratory spiking experiments that CWD surveillance via fecal sampling is possible from a range of mammalian and avian species. Such surveillance via fecal sampling could expand the toolkit of available approaches, offering a non-invasive alternative to current surveillance approaches, which typically requires animal capture. We then evaluated our methods using field-collected feces from CWD endemic areas, determining that this approach is able to detect PrP^CWD^ in the quantities present in coyote and cougar feces. Lack of tools that can be used to detect the presence of PrP^CWD^ in abiotic and biotic environmental samples has hampered our ability to address relevant ecological questions. This study directly addresses this need by developing an important surveillance tool for CWD, allowing investigation of the epidemiology of CWD at the community level.

The spiking experiments provided an assessment of how compatible RT-QuIC is with feces from different cervid consumers. Limits of relative PrP^CWD^ detection ranged from 10^5^ to 10 ng of spiked material, depending on species and spike dilution (Fig. 1, Fig. 2a, b). These dilutions of CWD-positive material are within the range of the relative PrP^CWD^ observed from feces of free-ranging coyote and cougar. Comparing among carnivore species, wolf and eagle samples had the most sensitive detection of brain derived PrP^CWD^ (Fig. 1), suggesting these species may be good early sentinels of CWD in an area. However, an assay cutoff time of 25 hours had to be applied to eagle results to remove false seeding events from unspiked samples, which could potentially result in loss of sensitivity. This was also the case for bear, crow, and raccoon, where assay cutoff times necessary to reduce false positives varied for each species. While these spiking experiments provide a baseline of compatibility for feces from a range of carnivores with the RT-QuIC assay, our study did not evaluate how intraspecific variation may affect the RT-QuIC assay. It is also worth noting that feces from animals in rehabilitation centers needed an assay cutoff time to reduce false seeding, whereas fecal samples from free-ranging animals did not require an assay cutoff time to avoid false positives. Thus, dietary variation or other species- or individual-specific differences may affect assay results. Further analyses of how differences in diet and fecal composition affect assay sensitivity and specificity may improve our understanding of assay strength and limitations.

Based on results of the spiking experiments, carnivore feces from certain cervid consumers could be a useful non-invasive approach to CWD surveillance, possibly able to detect CWD when prevalence rates in deer are very low. Our findings from both the Pine Ridge and Niobrara Valley DMUs highlight the usefulness of incorporating carnivore feces in CWD surveillance strategies. If predators select infected prey at a greater rate than uninfected prey, as has been observed with cougars^54^, predator feces may serve as early sentinels of CWD detection in areas where CWD is not yet endemic or where areas where surveillance efforts are limited, as we have demonstrated here with cougar feces. (Fig. 4, Table 3).

Differences in PrP^CWD^ recovery from feces spiked with CWD-positive WTD brain compared to the reference sample followed a similar trend among carnivores based on AFR values. Smaller differences were observed between the brain positive controls and spiked samples at the most and least concentrated spike dilutions compared to larger differences in detection at the mid-concentration dilutions (Fig. 2b). This hump-shaped variation in prion recovery across dilutions may be a result of the extraction process used for the feces samples following spiking, where some loss of seeding material likely occurred and was most sensitively observed at these spiking dilutions for each feces sample. It is also possible that the variability in AFRs that occurred for the 10^6^ ng and 10 ng concentrations of pure CWD-positive WTD BH was similar enough to that observed in many of the spiked fecal samples, that the differences were lower by comparison to the less variable AFRs from all other dilutions of pure CWD-positive WTD BH (Fig. 1, Fig. 2b). Additionally, the compounds that a given sample is composed of may influence the RT-QuIC reaction itself, as has been demonstrated with the inhibitory effect of certain sample types^55, 56^. Thus, it remains possible that this trend may be due to similar fecal biochemical compositions from each species that are resulting in similar recovery effects.

Detection of brain-derived PrP^CWD^ from wolf feces was the most sensitive across species, did not require an assay cutoff time to distinguish true seeding from false seeding events, and could be relatively easy to collect on a seasonal basis in more temperate CWD endemic areas such as the upper Midwest. The sensitivity and specificity results from the spiking assay and ease of sample collection suggest that wolf may be an optimal candidate for incorporating into CWD surveillance strategies, within its range. However, there are other considerations when designing fecal sampling programs in different environments, such as land access, effort required for sample collection, and species density. From the selection of species evaluated here, most of the mammal and avian species, with the exception of fox, may be candidates for further evaluation by RT-QuIC. Although spiking assays for coyote and cougar feces revealed moderate sensitivity compared to wolf (Fig 1), these species also did not require an assay cutoff time and exhibited 100% specificity, compared to ∼90% specificity for wolf (1/8 false seeding rate; Fig. 1), suggesting they may also be good candidates for CWD surveillance. Raven may also be a good candidate due to smaller differences in detection of PrP^CWD^ compared to the reference sample, however ease of collection of avian feces may limit utility for determining CWD occurrence in a given area compared with mammalian feces. In this study, we readily had access to coyote and cougar feces from areas where CWD is either endemic or has been detected at low prevalence. We demonstrated detection of PrP^CWD^ using spiking assays for these two species, yet they were not as sensitive as wolf samples. Because these larger scavenger and predator species tend to consume larger amounts of biomass in one feeding—and thus, presumably more PrP^CWD^— detection of PrP^CWD^ from their feces may be more likely than fecal samples from avian species. This suggests that analysis of mammalian carnivore feces may offer a way to advance CWD surveillance strategies.

Detection of PrP^CWD^ within some of the individual field collected Nebraska cougar feces subsamples was more variable than Wisconsin coyote subsamples (Fig. 3, Fig. 4, Table 2, Table 3). This may have been the result of differences in sample age, as increased sample age may reduce detection by RT-QuIC for some sample types^57^. Wisconsin coyote feces were less than a week old and had seeding ratios similar to the Pine Ridge 2021 cougar feces. Given that the more recently collected Nebraska cougar feces (Pine Ridge 2021) had higher overall AFRs in samples with PrP^CWD^ present and the most consistent detection among all other samples in that group compared to the other groups of Nebraska cougar feces (Niobrara 2012, 2014; Pine Ridge 2010), suggests that this variation could have been due to differences in sample age. In addition, lower amounts of seeding material have been shown to result in more variable AFRs using RT-QuIC^31^, thus it is also possible that this variability in more recent Nebraska cougar fecal subsamples could be the result of overall lower levels of PrP^CWD^ present in the sample compared to Wisconsin coyote samples. Homogenizing or mixing the whole sample prior to RT-QuIC might yield more consistent seeding from a given sample. However, because microparticles can lead to complications in the RT-QuIC^39^, we chose to initially evaluate subsampling of primary samples rather than whole sample-homogenization in this study. Future studies comparing homogenized vs. unhomogenized samples for a given species could help determine how homogenization of a whole sample influences RT-QuIC reactions, recovery of PrP^CWD^, and detection sensitivity and specificity. Additionally, factors such as amount of biomass consumed, gut residence time of PrP^CWD^ and time spent at carcass sites, or whether they are visiting a site repeatedly and feeding over the course of several days are factors that warrant further investigation.

The results of the Wisconsin field collected coyote feces found at deer mortality sites demonstrate that triplicate subsampling of each fecal sample was sufficient to overcome the variability of sensitivity for a given sample and demonstrate PrP^CWD^ presence using the RT-QuIC assay (Fig. 3, Table 2). Sample C3 had the most variable seeding activity and also had the lowest average AFRs compared to all other Wisconsin coyote samples, suggesting that the variability was likely due to lower overall levels of PrP^CWD^ present in this sample, a common response of the RT-QuIC assay to samples with lower prion loads^31^ (Fig. 3, Table 2). In addition, relative loads of PrP^CWD^ of each WI coyote fecal sample were reduced compared to tissues from the CWD-infected deer carcass they may have been scavenging (Fig. 3).

Because coyotes reportedly leave numerous scent marking feces within 1 to 80 m of the carrion they are consuming, we find it reasonable to consider that some of these fecal samples came from coyote that had scavenged on the respective carcass near the location their fecal samples were collected^49^. It is also worth considering that since one of the coyote fecal samples was collected on the periphery of a CWD-negative carcass site, —yet still had PrP^CWD^ present, some of these individuals may have been feeding on more than one carcass. This makes it difficult to interpret the reason for reduced loads of PrP^CWD^ in fecal samples compared to host tissue, to ascertain if reduced PrP^CWD^ loads are an effect of digestion or an effect of mixed consumption of different tissues from healthy and CWD-infected carcasses or other species in their diet. The reduced prion loads in coyote feces compared to deer tissue could be due to different or mixed dietary sources than just the sampled carcass, prion removal during digestion^27^, a species or sample type effect on prion recovery^58^, or a loss of seeding material during sample extraction. Additional studies assessing the relationship between CWD-infected biomass consumption and field-collected predator and scavenger fecal samples could help decipher the role of these species with respect to PrP^CWD^ environmental removal or deposition. Overall, these results support our hypothesis that CWD surveillance using carnivore feces may provide a useful estimate of CWD occurrence.

The developments we have reported here are critical steps in elucidating the roles of scavengers and predators in CWD epidemiology and for advancing and complimenting existing CWD surveillance strategies. Adding or altering ongoing surveillance efforts to include collection and analysis of predator and scavenger feces may call for additional funding, personnel, and logistical considerations by wildlife management agencies. However, CWD surveillance often relies upon hunter-harvest submission of tissues, followed by costly testing programs, typically funded by the state or province wildlife management agency. The ability to detect PrP^CWD^ in feces from cervid-consuming predators and scavengers may offer a cost-effective surveillance alternative for potentially early CWD detection and management action and could provide an efficient means to surveille at-risk areas neighboring CWD outbreak zones or areas where hunter-harvesting or hunter-submitted sampling is low, or where noninvasive approaches are desirable.

## ACKNOWLEDGEMENTS

This work was supported by the Wisconsin Department of Natural Resources (37000-0000009433 and 37000-0000010649). We thank the Yellowstone Biological Technicians that helped collect fecal samples for this project, especially Connor Meyer, Nikki Tatton, Jeremy SunderRaj, and Jack Rabe. Any use of trade, firm, or product names is for descriptive purposes only and does not imply endorsement by the U.S. Government.

## AUTHOR CONTRIBUTIONS

Conception and design of the work: H.N.I.; acquisition, analysis, and interpretation of the RT-QuIC data: H.N.I., W.C.T, S.S.L; writing the original draft: H.N.I., revision and edition of the final manuscript: H.N.I., E.E.B, S.P.W., M. H., D.R.S, K.W., D.P.W., T.N., D.J.S., S.S.L., and W.C.T. All authors approved and agreed to the submitted version of the present manuscript.

## DATA AVAILABILITY

All data generated or analyzed during this study are included in this published article.

## ADDITIONAL INFORMATION

Supplementary information is available for this paper. Correspondence and requests for materials should be addressed to

