## Supplementary Material for "Detection of prions from spiked and free-ranging carnivore feces"

**TABLES AND FIGURES**

**Supplementary Table S1.** Ordered differences of the Least Squares mean amyloid formation rates between all spiked feces and chronic wasting disease (CWD)-positive brain homogenate used for spiking fecal samples. Statistical significance was determined by Tukey HSD multiple comparisons test.

| **Feces type** | **Feces type comparison** | **Difference** | **Pooled standard error** | **Lower CL** | **Upper CL** | **p-value** |
| --- | --- | --- | --- | --- | --- | --- |
| Raven | Fox | 0.031677 | 0.003655 | 0.020052 | 0.043301 | 0.00E+00 |
| Raven | Cougar | 0.030095 | 0.003655 | 0.018471 | 0.04172 | 0.00E+00 |
| Raven | Raccoon | 0.011151 | 0.003655 | -0.00047 | 0.022775 | 7.25E-02 |
| Raven | Coyote | 0.010118 | 0.003655 | -0.00151 | 0.021743 | 1.51E-01 |
| Raven | Bear | 0.008386 | 0.003655 | -0.00324 | 0.020011 | 3.95E-01 |
| Raven | Wolf | 0.003546 | 0.003655 | -0.00808 | 0.015171 | 9.94E-01 |
| Cougar | Fox | 0.001582 | 0.003655 | -0.01004 | 0.013206 | 1.00E+00 |
| Wolf | Fox | 0.028131 | 0.003655 | 0.016506 | 0.039755 | 0.00E+00 |
| Wolf | Cougar | 0.026549 | 0.003655 | 0.014925 | 0.038174 | 2.22E-11 |
| Wolf | Raccoon | 0.007605 | 0.003655 | -0.00402 | 0.019229 | 5.42E-01 |
| Wolf | Coyote | 0.006572 | 0.003655 | -0.00505 | 0.018197 | 7.36E-01 |
| Wolf | Bear | 0.00484 | 0.003655 | -0.00678 | 0.016465 | 9.48E-01 |
| Wolf | Eagle | 0.021598 | 0.003655 | 0.009973 | 0.033222 | 3.17E-07 |
| Eagle | Fox | 0.049729 | 0.003655 | 0.038104 | 0.061353 | 0.00E+00 |
| Eagle | Cougar | 0.048147 | 0.003655 | 0.036523 | 0.059772 | 0.00E+00 |
| Eagle | Raven | 0.018052 | 0.003655 | 0.006427 | 0.029676 | 4.94E-05 |
| Eagle | Raccoon | 0.029202 | 0.003655 | 0.017578 | 0.040827 | 0.00E+00 |
| Eagle | Coyote | 0.02817 | 0.003655 | 0.016545 | 0.039794 | 0.00E+00 |
| Eagle | Bear | 0.026438 | 0.003655 | 0.014813 | 0.038062 | 4.05E-11 |
| Bear | Fox | 0.023291 | 0.003655 | 0.011666 | 0.034915 | 2.18E-08 |
| Bear | Cougar | 0.021709 | 0.003655 | 0.010085 | 0.033334 | 2.67E-07 |
| Bear | Raccoon | 0.002765 | 0.003655 | -0.00886 | 0.014389 | 9.99E-01 |
| Bear | Coyote | 0.001732 | 0.003655 | -0.00989 | 0.013357 | 1.00E+00 |
| Raccoon | Fox | 0.020526 | 0.003655 | 0.008902 | 0.032151 | 1.58E-06 |
| Raccoon | Cougar | 0.018945 | 0.003655 | 0.00732 | 0.030569 | 1.49E-05 |
| Crow | Fox | 0.065923 | 0.003655 | 0.054299 | 0.077548 | 0.00E+00 |
| Crow | Cougar | 0.064342 | 0.003655 | 0.052717 | 0.075966 | 0.00E+00 |
| Crow | Raven | 0.034246 | 0.003655 | 0.022622 | 0.045871 | 0.00E+00 |
| Crow | Raccoon | 0.045397 | 0.003655 | 0.033773 | 0.057022 | 0.00E+00 |
| Crow | Coyote | 0.044365 | 0.003655 | 0.03274 | 0.055989 | 0.00E+00 |
| Crow | Bear | 0.042633 | 0.003655 | 0.031008 | 0.054257 | 0.00E+00 |
| Crow | Wolf | 0.037792 | 0.003655 | 0.026168 | 0.049417 | 0.00E+00 |
| Crow | Eagle | 0.016195 | 0.003655 | 0.00457 | 0.027819 | 5.03E-04 |
| Coyote | Fox | 0.021559 | 0.003655 | 0.009934 | 0.033183 | 3.36E-07 |
| Coyote | Cougar | 0.019977 | 0.003655 | 0.008353 | 0.031602 | 3.50E-06 |
| Coyote | Raccoon | 0.001033 | 0.003655 | -0.01059 | 0.012657 | 1.00E+00 |
| Brain | Fox | 0.107407 | 0.003655 | 0.095782 | 0.119031 | 0.00E+00 |
| Brain | Cougar | 0.105825 | 0.003655 | 0.094201 | 0.11745 | 0.00E+00 |
| Brain | Raven | 0.07573 | 0.003655 | 0.064105 | 0.087354 | 0.00E+00 |
| Brain | Raccoon | 0.086881 | 0.003655 | 0.075256 | 0.098505 | 0.00E+00 |
| Brain | Coyote | 0.085848 | 0.003655 | 0.074223 | 0.097472 | 0.00E+00 |
| Brain | Bear | 0.084116 | 0.003655 | 0.072491 | 0.09574 | 0.00E+00 |
| Brain | Wolf | 0.079276 | 0.003655 | 0.067651 | 0.0909 | 0.00E+00 |
| Brain | Eagle | 0.057678 | 0.003655 | 0.046054 | 0.069303 | 0.00E+00 |
| Brain | Crow | 0.041483 | 0.003655 | 0.029859 | 0.053108 | 0.00E+00 |

**Supplementary Table S2.** Percent recovery of seeding material based on amyloid formation rate (AFR) values of 10-fold dilution spikes in fecal samples for each species to AFR values of 10-fold dilutions of the reference sample.

| **Species** | **Spike dilutions and percent recovery based on AFR** | | | | | |
| --- | --- | --- | --- | --- | --- | --- |
|  | 10^6^ | 10^5^ | 10^4^ | 10^3^ | 100 | 10 |
| Crow | 96% | 94% | 78% | 45% | 0% | 0% |
| Eagle | 62% | 59% | 57% | 62% | 43% | 0% |
| Wolf | 68% | 47% | 32% | 33% | 44% | 60% |
| Raven | 66% | 45% | 43% | 37% | 0% | 0% |
| Bear | 50% | 42% | 38% | 15% | 10% | 0% |
| Coyote | 68% | 42% | 27% | 3% | 8% | 0% |
| Raccoon | 55% | 48% | 29% | 12% | 0% | 0% |
| Cougar | 39% | 15% | 7% | 8% | 0% | 0% |
| Fox | 36% | 12% | 5% | 5% | 0% | 13% |

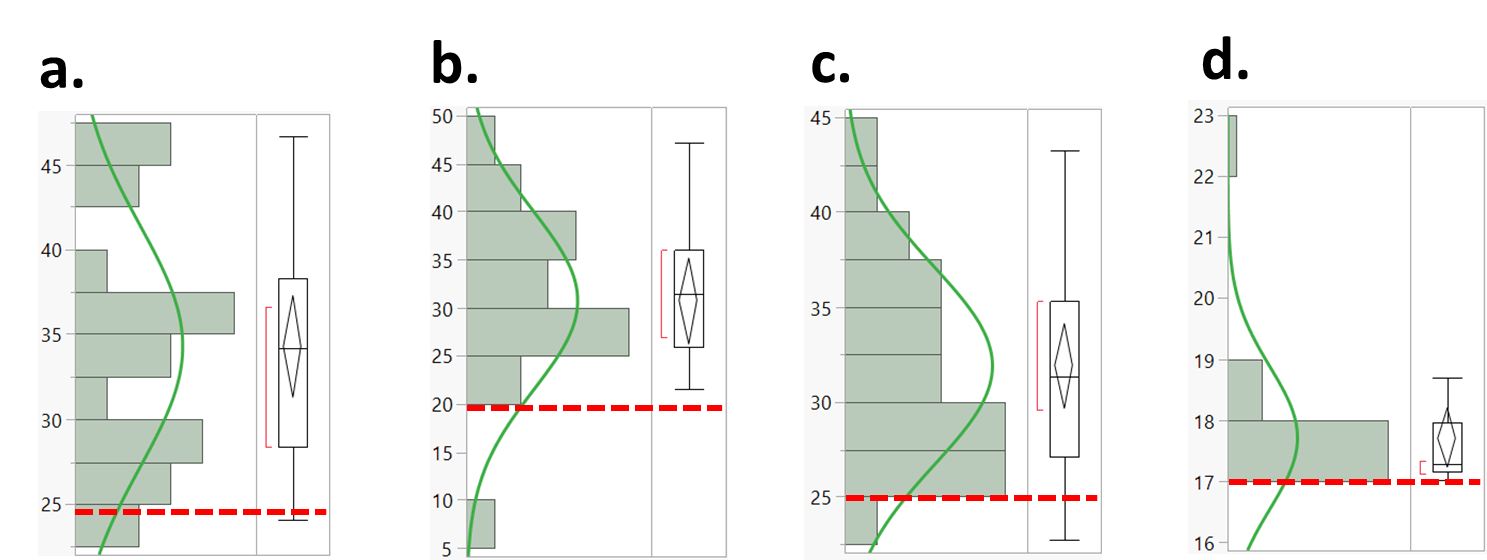

**Supplementary Figure S1** Histogram plots depicting empirical distributions of threshold times for 24 technical replicates of unspiked feces from **(a)** bear (34.3, ±7.1), **(b)** crow (M,30.7; SD, 9.2), **(c)** eagle (M, 31.9; SD, 5.2), and **(d)** raccoon (M, 17.7; SD, 1.2). Horizontal dotted red lines indicate cycle end-times that excluded ≥95% of the false seeding that occurred for each of these feces.

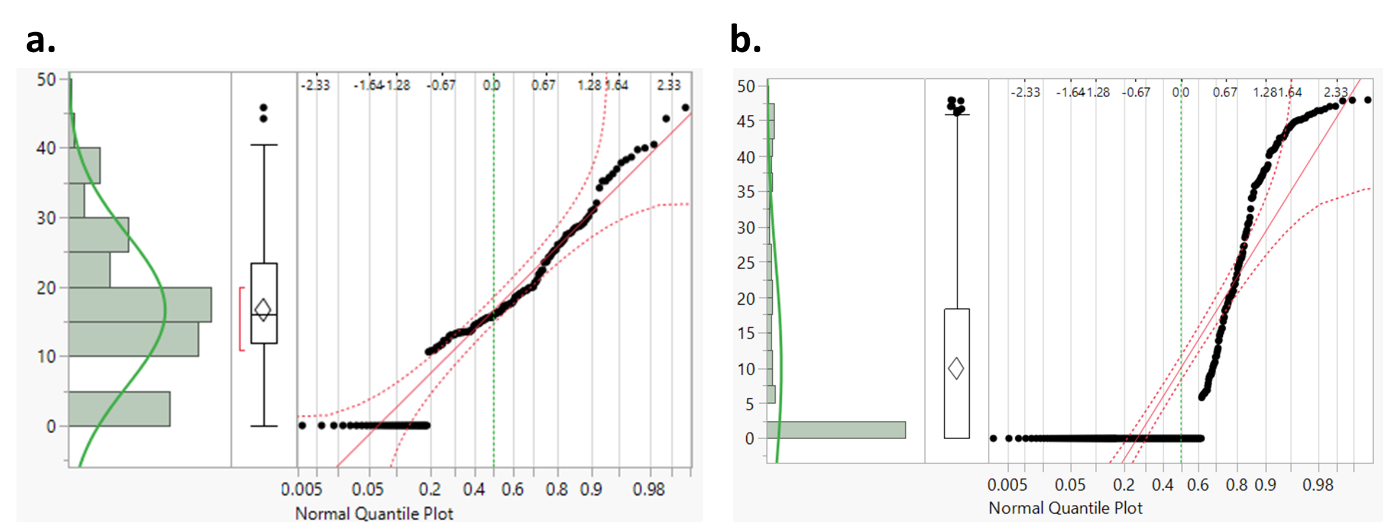

**Supplementary Figure S2.** Non-normal distributions of the time (hours)-to-threshold data for predator and scavenger fecal samples. Histogram and normal quantile plot depicting distribution of hours to threshold for (a) coyote fecal sample data depicted in “Figure 3” (N, 168; M, 16.6; SD, 10.9; Goodness of Fit P, < .0001) and (b) cougar fecal sample data depicted in “Figure 4” (N, 384; M, 9.9; SD, 15.27, Goodness of Fit P, < .0001)
